# An mRNA silencing mechanism reliant on the cooperation between REGE-1/Regnase-1 and RLE-1/Roquin-1

**DOI:** 10.1101/2021.12.13.472375

**Authors:** Daria Sobańska, Alicja A Komur, Agnieszka Chabowska-Kita, Julita Gumna, Pooja Kumari, Katarzyna Pachulska-Wieczorek, Rafal Ciosk

## Abstract

Regnase-1 is an evolutionarily conserved endoribonuclease, degrading diverse mRNAs important, among others, for immune homeostasis, development, and cancer. There are two competing models of Regnase-1 mediated mRNA silencing. One model postulates that Regnase-1 works together with another RNA-binding protein, Roquin-1. The other model proposes that the two proteins function separately. Studying the *C. elegans* Regnase-1 ortholog, REGE-1, we have uncovered a functional relationship between REGE-1 and the nematode counterpart of Roquin-1, RLE-1. While REGE-1 and RLE-1 associate with mRNA independently of each other, both proteins are essential for mRNA silencing. Intriguingly, the functional interdependence between REGE-1 and RLE-1 is reminiscent of the proposed cooperation between mammalian Regnase-1 and Roquin-1, which may underlie a prototypic silencing mechanism involving both proteins.

## INTRODUCTION

mRNAs are regulated at many levels, including at the level of transcript stability. The mRNA specificity is mediated by *cis*-acting RNA motifs, which are associated with diverse RNA binding proteins (RBPs) and/or non-coding RNAs. An important class of RBPs affecting RNA stability are ribonucleases (RNases). RNases degrade RNA either from its ends (exoribonucleases) or via internal cleavages (endoribonucleases). A well-known example of endoribonucleases is Regnase-1 (regulatory RNase 1), also known as Mcpip1 or Zc3h12a (1,2). Regnase-1 has a critical role in mammalian innate immunity, as well as in cancer, infection, angiogenesis, cell differentiation, and apoptosis (3). Regnase-1 was initially reported to co-regulate mRNAs with another RBP, Roquin-1, which was suggested to recruit Regnase-1 to specific mRNA targets during T-cell activation (4). However, more recent studies, in various cell types, suggest that Roquin-1 and Regnase-1, although controlling overlapping mRNAs, function in different subcellular compartments and by entirely different molecular mechanisms (5,6). According to the latter, independent model, Regnase-1 degrades translationally active mRNAs with the help of Upf1 (a helicase known for its role in nonsense-mediated RNA decay, NMD) (6), which facilitates Regnase-1 mediated endonucleolytic cleavage by unwinding structured RNA (7). By contrast, Roquin-1 was shown to target translationally inactive mRNAs, promoting their exonucleolytic degradation by recruiting the CCR4-NOT deadenylase complex (5,8). In addition to promoting mRNA decay, Roquin-1 was also reported to inhibit mRNA translation (9).

Both proteins bind the 3’ untranslated regions (3’UTRs) of target mRNAs (5). In RNA cleavage assays, Regnase-1 did not display a strong preference for RNA sequences or structures (10). Other studies reported that Regnase-1 targets harbor stem-loops (SLs), and mutations disrupting these SLs abolish Regnase-1-mediated degradation (10). Whether these SLs recruit Regnase-1 remains, however, unclear. Finally, according to the co-regulation model, the RNA specificity of Regnase-1 comes from Roquin-1, which, by binding specific RNA elements, recruits Regnase-1 for degradation (4). However, there is currently no evidence for a physical association between Regnase-1 and Roquin-1. By contrast, the RNA specificity of Roquin-1, as well as the related proteins collectively referred to as “Roquin”, is well understood. Roquin binds specific SLs, with the loops containing conserved motifs of either three nucleotides (called constitutive decay elements, CDEs) (8), or six nucleotides (alternative decay elements, ADEs) (11,12). The binding to these SLs is mediated by the so-called ROQ domain (11,13).

Recently, we characterized the *Caenorhabditis elegans* ortholog of Regnase-1, which we called REGE-1 (REGnasE-1) (14). Studying its biological functions, we identified a key target of REGE-1, the *ets-4* mRNA, encoding a conserved transcription factor regulating body fat and cold resistance. REGE-1 shares the domain organization with Regnase-1, and its RNase activity is essential for *ets-4* mRNA degradation. Within the *ets-4* mRNA 3’UTR is a short fragment, 115 nucleotide-long, which is sufficient for REGE-1 mediated mRNA silencing (REGE-1 cleaves RNA within that fragment). However, the specificity of REGE-1 towards *ets-4* and the potential involvement of other players remained unknown. The independent model of Regnase-1 mediated silencing is substantiated by a greater body of evidence, and the *C. elegans* REGE-1 and RLE-1 are highly similar to their human counterparts (15). Thus, one might have anticipated independent functions of the nematode proteins. Instead, we demonstrate that REGE-1 and RLE-1 are functionally connected. Dissecting the RNA features underlying the regulation by both proteins, we find that, even though the proteins bind mRNA separately, they cooperate in inducing mRNA decay, similar to previously proposed co-regulation by Regnase-1 and Roquin-1 (4). By demonstrating a conserved functional relationship between REGE-1/Regnase-1 and RLE-1/Roquin, our work provides a unique perspective on the evolution of a posttranscriptional regulatory mechanism, playing important functions in health and disease.

## MATERIALS AND METHODS

### Biological resources

All C. *elegans* strains used in these studies are listed in Table S1.

HEK 293T cells were ordered from ATCC, USA (https://www.atcc.org/products/crl-3216).

### *C. elegans* handling and genetic manipulation

Animals were grown at 20°C on standard NGM plates, fed with the OP50 *E. coli* bacteria (16). The CRISPR/Cas9 genome editing was performed by SunyBiotech, China, to generate the *rle-1* ROQ domain mutant (allele *syb517*) and to C-terminally tag *rle-1* with GFP-FLAG (allele *syb1279*). The strain expressing F1SΔRCE reporter (allele *sybSi111*) was generated (by SunyBiotech) using the MosSCI method, utilizing the insertion locus ttTi5605.

RNAi of individual genes was performed by feeding animals with bacteria expressing double-stranded RNA, beginning from the L1 larval stage until the young adult stage at 20°C, except the RNAi of CCR4-NOT deadenylase complex that was done from the L4 stage. The L4440 (empty) vector was used as a negative RNAi control. The RNAi clones used in this study came from either Ahringer or Vidal libraries.

### Quantification of the *ets-4* 3’ UTR GFP reporter

Images for quantification of GFP intensity of reporter strains were acquired with an Axio Imager.Z2 (Carl Zeiss, Germany), equipped with Axiocam 506 mono digital camera (Carl Zeiss, Germany), and a Plan-Apochromat 63x/1.40 Oil DIC M27 objective. Images, acquired with the same camera settings, were processed with ZEN 2.5 (blue edition) microscope software in an identical manner. The signal intensity of a circular area of 150-pixels diameter of five to seven gut nuclei from five animals per condition was measured in ImageJ (17) and normalized to the background. In addition, 30–35 animals per strain were visually inspected for GFP expression. Statistical analysis on all of the experiments was performed using GraphPad Prism 8. The statistical method used to calculate P-value is indicated in the figure legends.

### qRT-PCR

Around 1000 young adult *C. elegans* were collected at 20°C, washed 2 times in M9 buffer, and flash-frozen in Trizol. Total RNA was isolated using Direct-zol RNA Miniprep Kit (Zymo Research, USA, Cat. No. R2053). 2000 ng of RNA was used to prepare cDNA with High-Capacity cDNA Reverse Transcription Kit (Applied Biosystems, USA, Cat. No. 4368814). cDNA was diluted at 1:10 and 4 µl of template cDNA was mixed with AMPLIFY ME SG Universal Mix (Blirt, Poland, Cat. No. AM02-200). Ct values were calculated using Light Cycler 480 (Roche, Switzerland). The *tbb-2* (beta-tubulin) was used as the reference gene. Statistical analysis on all of the experiments was performed using GraphPad Prism 8. The statistical method used to calculate P-value is indicated in the figure legends. qRT-PCR primers used in this study are listed in Table S2.

### The assay for *C. elegans* cold survival

Cold survival experiments were performed as published (14). Specifically, prior to cold adaptation, animals were grown at 20°C for two generations on OP50. They were then synchronized by bleaching, and L1 larvae were grown until day 1 of adulthood at 20°C. On day 1 of adulthood, they were cold-adapted at 10°C for 2 hrs and then shifted to 4°C. Animals were sampled at indicated intervals, and their survival was scored after 24 h recovery at 20°C.

### Oil Red O staining

Oil red O staining was performed as published (18). 0.5 g of Oil Red O was mixed with 100 ml isopropanol and stirred for 24 h, protected from direct light. This solution was diluted in water to 60%, stirred for 12 h, and filtered by 0.22 µm pore filter. About 3000 young adult animals were washed from the plates, washed three times with M9, and fixed with 75% isopropanol for 15 min with shaking at 1400 rpm. After fixation followed by spinning and removal of isopropanol, animals were suspended in 1 ml of 60% Oil red O and stained for 3 h on a shaker with maximum speed, covered with aluminium foil. After staining worms were washed four times with PBS-T. Stained animals were placed on 3% agar pads and imaged on Nikon SMZ25 with DeltaPix color camera with 60x zoom. All image-processing steps were done with the Fiji/ImageJ software (19). The calculations were made in two steps. The first step was to measure the signal from the red color. After conversion from RGB to HSB color space and background subtraction, red pixels were selected by color thresholding. A binary mask was created with the saturation channel and applied to the thresholded image. After conversion to 32-bit, zero pixel values were replaced by NaN. The integrated density of all remaining pixels was used as an index of the amount of red staining in the animals. In the next step, the area of the worm was calculated. The image was converted to 8-bits, background subtraction was made and a threshold was set to measure the surface of the particles. The signal from red pixels was compared to the worm’s area. 30 animals were imaged per strain and biological replicate. A two-tailed t-test was used to calculate significance with Graph Pad/Prism 8.

### Dual-luciferase assay

The 3’ UTRs were cloned into psiCheck-2 plasmid (Promega, USA, Cat. No. C8021) using the XhoI and NotI sites. Short fragment of *ets-4* 3’ UTR (F1S) was inserted into AscI site in the *unc-54* 3’ UTR, using Gibson assembly with Gibson assembly Master MIX (NEB, USA, Cat. No. E2611S). Primers used for fragment amplification are listed in Table S2. pFLAG-CMV2-Regnase-1 and pFLAG-CMV2-Regnase-1 D141N plasmids were kind gifts from Osamu Takeuchi. REGE-1 cDNA was cloned into pFLAG-CMV2 plasmid using NotI and KpnI restriction sites. HEK-293T cells were grown in Dulbecco’s modified Eagle medium without glucose supplemented with 10% heat-inactivated fetal calf serum. 10 ng of pFLAG expression vectors or empty plasmids and 50 ng psiCheck-2 plasmid were transfected using Effectene transfection reagent (Qiagen, Netherlands, Cat. No. 301425), following the manufacturer’s instructions. Transfection was done in at least three independent biological replicates of cells. Luciferase assay was done using a dual-luciferase reporter assay system (Promega, USA, Cat. No. E1910), following the manufacturer’s instructions. For each measurement, an average of two technical replicates was used for further analysis. Statistical analysis on all of the experiments was performed using GraphPad Prism 8. The statistical method used to calculate P-value is indicated in the figure legends.

### Western blot analysis

Western blot analysis was conducted as described previously (20). Primary antibodies diluted in 5% milk/PBS-Tween were: mouse anti-Actin (1:2000; Merck, Germany, Cat. No. MAB1501), rat anti-MYC (1:1000; Chromotek, Germany, Cat. No. 9e1-20) and rabbit anti-REGE-1 polyclonal antibody raised against the first 119 amino acids (1:1000; SDIX, USA). Detection was performed with horseradish peroxidase-conjugated secondary antibodies: horse anti-mouse (1:5000; Cell Signaling, USA, Cat. No. Cat. No. 7076S), goat anti-rat (1:5000; Cell Signaling, USA, Cat. No. 7077S), and goat anti-rabbit (1:5000; Cell Signaling, USA, Cat. No. 7074S), radiance ECL detection reagent (Azure Biosystems, Germany, Cat. No. AC2204) and c600 imaging system (Azure Biosystems, Germany). Western blotting was performed for all the biological replicates. One representative blot is shown in the results.

### RNA co-immunoprecipitation (RIP)

RNA co-immunoprecipitation (RIP) was performed on total lysates from *rege-1(rrr13)* or *rege-1(rrr13); rle-1(rrr44)* animals expressing a single copy rescuing RNase-dead REGE-1::GFP transgene. Worms were synchronized, grown at 20°C on standard NGM plates, and fed with the OP50 *E. coli* bacteria until the young adult stage. Pellets were prepared by harvesting worms in M9 buffer, washing by harvest buffer (100 mM KCl, 0.1% Triton X-100), and freezing in liquid nitrogen. Lysates from four biological replicates were prepared by grinding the frozen worm pellets using mortar and pestle, and dissolving them in the lysis buffer (50 mM Hepes pH 7.5, 150 mM KCl, 5 mM MgCl2, 0.1 % Triton X-100, 10% glycerol) supplemented with 150 mM PMSF, Complete EDTA-free protease inhibitors (Roche, Switzerland, Cat. No. 11873580001), 1 µM Pepstatin A, 0.3 µM Aprotinin, and 200 U RNase inhibitor - RNasin (Promega, Germany, Cat. No. N2511) at 4 °C. Lysates were cleared by centrifugation at 15,000 × g for 20 min at 4 °C and passed through a 0.45 µm syringe filter. IPs were performed by incubating lysates equivalent to 20 mg total protein supplemented with RNase inhibitor - RNasin (Promega, Germany, Cat. No. N2511) and 1 mM DTT with 80 μl of anti-GFP magnetic beads GFP-Trap (Chromotek, Germany, Cat. No. gtma-20) for 3h at 4°C while rotating. Protein extract was saved as input for the western blot (1/200), and for RNA isolation (1/20). After IP beads were washed three times with the lysis buffer. Half of the beads were resuspended in a 2x SDS sample loading buffer, boiled for 5 min at 90^°^C, and analyzed by western blot as described above. The other half of the beads were resuspended in 700 µL Trizol for RNA extraction and frozen in liquid nitrogen. RNAs were extracted using 70% chloroform and precipitated. Then samples were centrifuged, and pellets were washed with 70% ethanol, dried, and resuspended in RNase-free water. cDNAs were synthetized using QuantiTect Reverse Transcription Kit (Qiagen, Netherlands, Cat. No 205311). RT-qPCR was performed as described above. After RNA extraction and subsequent qPCR analysis, fold enrichment was calculated as previously described (14).

### RNA synthesis and modification

RNAs for RNA pulldown experiment were *in vitro* transcribed using MEGAshortscript kit (Thermo Fisher Scientific, USA, Cat. No. AM1354) according to the manufacturer’s protocol. The DNA templates for transcription were obtained by PCR amplification of the psiCheck-2 vectors using a forward primer containing an SP6 promoter sequence by Phusion HF polymerase (NEB, USA, Cat. No. M0530S). After transcription, RNAs were purified using RNeasy MinElute Cleanup Kit (Qiagen, Netherlands, Cat. No. 74204) according to the manufacturer’s protocol. RNA purity was confirmed on agarose gel in denaturing conditions. 50 pmol RNAs were biotinylated using Pierce RNA 3’ End Biotinylation Kit (Thermo Fisher Scientific, USA, Cat. No. 20160) according to the manufacturer’s protocol, extracted by chloroform: isoamyl alcohol, precipitated and resuspended in RNase-free water.

RNAs for the SHAPE experiment were *in vitro* transcribed using MAXIscript SP6 (Thermo Fisher Scientific, USA, Cat. No. AM1310M) according to the manufacturer’s protocol. Templates for *in vitro* transcription were obtained by PCR amplification of *ets-4* 3’UTR variants from “*pDONR P2R--P3 ets-4 3’UTR F1”* plasmid using a forward primer containing an SP6 promoter sequence. RNAs were DNase I-treated and recovered by EtOH precipitation. For RNA modification, 20 pmols of RNA were refolded in 20μl of buffer containing 10 mM Tris-HCl, pH 7.5, 100 mM KCl and 0.1 mM EDTA by heating 3 min at 95°C, slow cooling to 4°C, adding 100 μl H2O and 50 μl of 5x folding buffer (1x: 40 mM Tris-HCl, pH 7.5, 140 mM KCl, 0.2 mM EDTA, 5 mM MgCl2), and incubating for 10 min at 37°C. Folded RNA was divided equally into two tubes and treated with either 8 μl of 100 mM 1M7 in DMSO (reaction, (+)) or DMSO alone (control (-)) and allowed to react for 5 min at 37°C. RNA was recovered using Direct-zol RNA MiniPrep Kit (Zymo Research, USA, Cat. No. R2050) and resuspended in 25 μl of H2O.

### *In vitro* translation

DNA templates for *in vitro* translation were obtained by cloning RLE-1 and REGE-1 RNase-dead mutant cDNAs with N-terminal FLAG tag and C-terminal MYC tag into pCS2+ plasmid using ClaI and XbaI restriction sites using primers listed in Table S2. Proteins were *in vitro* translated using TnT Quick Coupled Transcription/Translation System (Promega, USA, Cat. No. L1170) according to the manufacturer’s protocol.

### RNA pulldown

BioMag Nuclease-Free Streptavidin particles (Bangs Laboratories, USA, Cat. No. BM568) were washed and equilibrated with the binding buffer (5 mM HEPES pH 7.5, 70 mM KCl, 20 mM MgCl2, 1 mM DTT, 0.1% Triton X-100, 1% glycerol, 20 µg/ml yeast tRNA). 7 pmol biotinylated RNAs were incubated with streptavidin beads in the binding buffer supplemented with 80 U RNasin (Promega, Germany, Cat. No. N2511) and 50 µg yeast tRNA (Thermo Fisher Scientific, USA, AM7119) for 2 h at 4^°^C with shaking. Unbound RNAs were washed with the binding buffer, beads were resuspended in the binding buffer, and incubated with reticulocyte lysates with *in vitro* translated proteins for 2 h at 4^°^C with shaking. After incubation, streptavidin beads were washed, resuspended in a 2x SDS sample loading buffer, and proteins were separated using SDS-PAGE. Proteins bound to RNAs were detected based on their size using western blot analysis as described above.

### SHAPE structure probing

Detection of 2’-O-adducts was performed using reverse transcription with [^32^P]-labeled primers. Briefly, 1 μl of primer (AGGAATATGTTCTACAACGAACAGT) was added to the 0.5 pmol of (-) and (+) RNA, and 12 μl of primer-template solutions were incubated at 95°C for 2 min, followed by 60°C for 5 min, and 52°C for 2 min. RNA was reverse transcribed at 52°C for 10 min by Superscript III (Thermo Fisher Scientific, USA, 12574026). Samples and sequencing ladders were purified by EtOH precipitation. Primer extension products were separated on denaturing PAA (8M urea). The gels were quantitatively analyzed by phosphoimaging using FLA-5100 phosphoimager (Fujifilm, Japan) with MultiGaugeV 3.0 software.

## RESULTS

### REGE-1 and Regnase-1 are functional homologs

Recently, we characterized the *C. elegans* ortholog of Regnase-1, REGE-1, which forms an auto-regulatory module with its key target, the ETS-4 transcription factor (14). REGE-1 controls ETS-4 levels via the degradation of *ets-4* mRNA, induced by REGE-1 mediated cleavage within the *ets-4* 3’ UTR (14). By regulating the levels of ETS-4, REGE-1 controls various aspects of animal physiology, including fat metabolism and cold resistance (Fig. 1A) (14,21). The *C. elegans* REGE-1 and mammalian Regnase-1 share the same domain organization, suggesting their functional conservation (22). To test it, we examined, in mammalian cells, whether REGE-1 can repress human mRNAs and Regnase-1 can repress nematode mRNA targets. Specifically, we transfected human embryonic kidney (HEK293T) cells with the so-called dual-luciferase (firefly/*Renilla*) reporter constructs, where one reporter is used as a reference and the other as a query, containing 3’ UTR sequence from a target mRNA. We tested reporters carrying 3’ UTRs of two Regnase-1 targets, *mIL6* (4,5) and *mOX40* (11,23), as well as the 3’ UTR of nematode *unc-54* mRNA that served as a negative control (Fig. 1B). Additionally, we tested the silencing of *ets-4* mRNA, which, as we previously showed, is mediated by a short fragment of the *ets-4* 3’UTR dubbed F1S. When “transplanted” into (otherwise unregulated) *unc-54* 3’ UTR, the F1S fragment is sufficient for REGE-1 mediated silencing (14). We co-expressed the above reporter constructs with REGE-1 or Regnase-1, or the RNase-dead variant Regnase-1 D141N (Fig. 1B-C). We found that, when expressed in mammalian cells, REGE-1 and Regnase-1 were interchangeable; while REGE-1 repressed the known targets of Regnase-1 (*IL6* and *OX40*), Regnase-1 repressed the REGE-1 targets (*ets-4* and *unc-54* carrying the F1S fragment) (Fig. 1B-C). Expectedly, the RNase activity of Regnase-1 was crucial for the silencing of REGE-1 targets, as the RNase-dead variant of Regnase-1 was unable to induce the silencing (Fig. 1C). Additionally, we saw a protective effect of Regnase-1 D141N mutant on *ets-4* mRNA decay (Fig. 1C), which was similar to previous observations on mammalian *IL6* (5).

**Figure 1.**
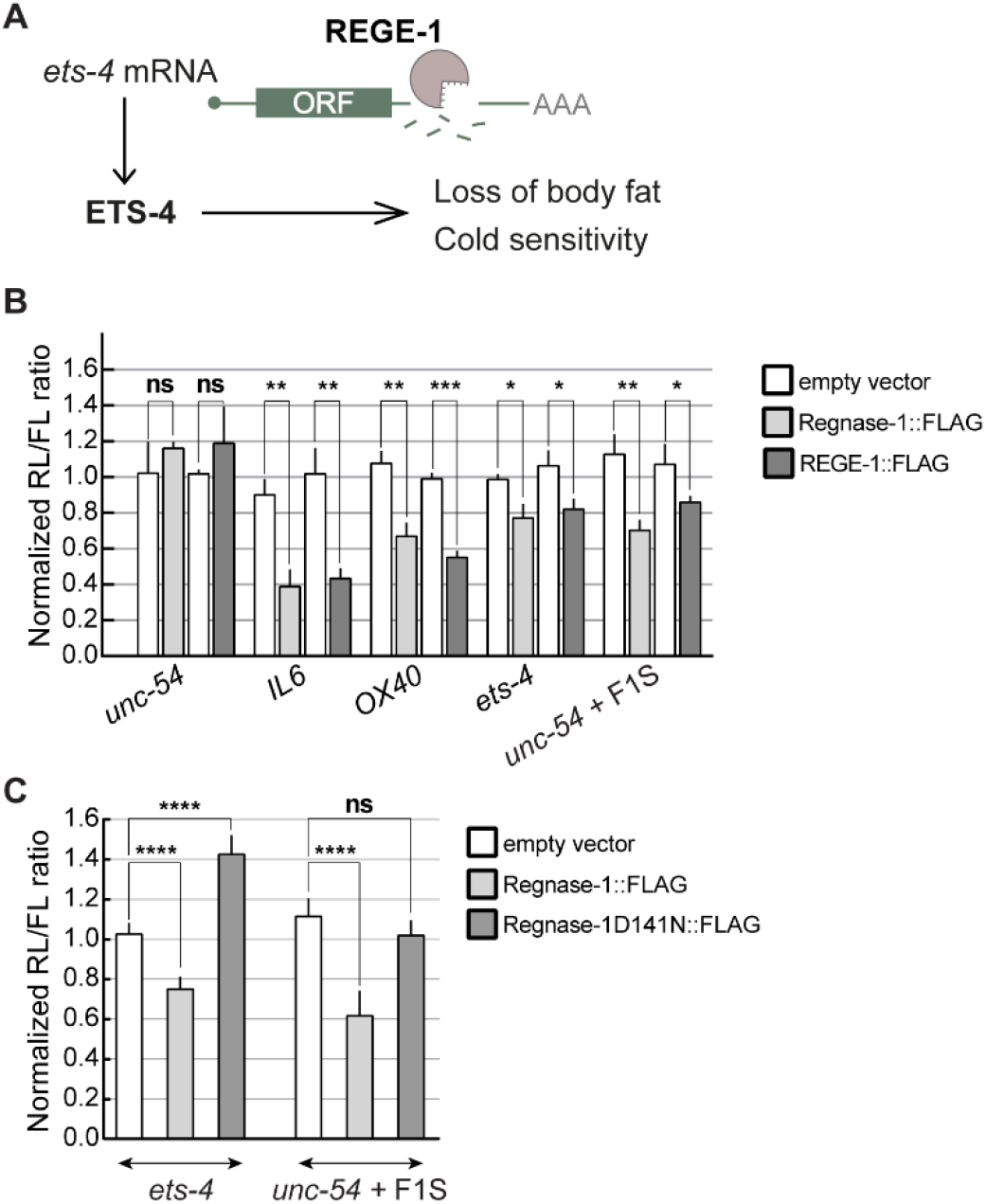
REGE-1 and Regnase-1 are functional homologs. (A) A model for REGE-1 mediated control of fat metabolism and cold resistance. By degrading the *ets-4* mRNA, REGE-1 lowers the levels of transcription factor ETS-4. This leads to altered expression of ETS-4 target genes, affecting the accumulation of body fat and cold resistance (14). (B) In HEK cells, REGE-1 and Regnase-1 are interchangeable and sufficient for mRNA silencing. HEK cells were transfected with reporters expressed under the control of various 3’ UTRs: from Regnase-1 targets (*OX40, IL6*), REGE-1 targets (*ets-4* and *unc-54* + F1S), and from an unregulated *C. elegans* mRNA (*unc-54*). Additionally, cells were co-transfected with constructs expressing empty vector (white bars), or mouse Regnase-1 (light grey bars), or worm REGE-1 (dark grey bars). Expression of the *Renilla* luciferase was controlled by the indicated 3’ UTRs and normalized to the firefly luciferase expression. Two-tailed P-values were calculated with an unpaired Student t-test, using empty vector results as a reference. Bars represent the mean value from three independent biological replicates (N = 3). Error bars represent SEM; * indicates P < 0.05, ** indicates P < 0.001, *** indicates P = 0.0001, “ns” = not significant. (C) The RNase activity of Regnase-1 is crucial for the silencing of REGE-1 targets. Same as in B, except that HEK cells were co-transfected with reporters controlled by 3’ UTRs of REGE-1 targets (*ets-4* and F1S in *unc-54*), and additionally constructs expressing empty vector (white bars), mouse Regnase-1 (light grey bars), or its RNase-dead mutant (Regnase-1 D141N; dark grey bars). Two-tailed P-values were calculated using unpaired Student t-test using empty vector results as a reference. Bars represent the mean value from five independent biological replicates (N = 5). Error bars represent SEM; **** indicates P < 0.0001, “ns” = not significant.

### The REGE-1 mediated mRNA silencing requires RLE-1, the *C. elegans* counterpart of Roquin-1

Considering the functional similarity between REGE-1 and Regnase-1, we asked whether REGE-1 uses one of the mechanisms proposed for Regnase-1. We thus tested the possible involvement of SMG-2 (the homolog of Upf1, reported to function with Regnase-1 according to the independent model) and RLE-1 (the *C. elegans* counterpart of Roquin-1, reported to function either independently or together with Regnase-1) studying mutant and wild-type *C. elegans* strains (Fig. 2A). While the expression level of *ets-4* mRNA in *smg-2(-)* mutants was comparable to the wild type, it increased in *rle-1(-)* mutants (Fig. 2B). Moreover, the magnitude of this increase was similar to that observed in either *rege-1(-)* single, or *rege-1(-); rle-1(-)* double mutants (Fig. 2B). Finally, underscoring the functional connection between both proteins, the RLE-1 dependent repression of *ets-4* was mediated, similar to REGE-1, by the *ets-4* 3’ UTR (Fig. 2C). Importantly, the loss of *ets-4* silencing, observed in *rle-1(-)* or *rle-1(RNAi)* animals, was not caused by the loss of REGE-1. Quite the opposite, in RLE-1 depleted animals, we observed increased levels of *rege-1* mRNA and REGE-1 protein (Fig S1A), which could be explained by ETS-4 mediated upregulation of *rege-1* transcription (14). At the same time, the expression of RLE-1, which, in agreement with previous studies, is enriched in the intestine (15), was not affected by the depletion of REGE-1 (Fig. S1B). To summarize, while SMG-2/Upf1 does not seem to be involved in REGE-1 mediated silencing, RLE-1 is as important for the silencing as is REGE-1. When functioning independently of Regnase-1 (independent model), Roquin was shown to repress mRNAs by recruiting the CCR4-NOT deadenylase complex (5,24). This is why we additionally examined the potential involvement of components of the nematode CCR4-NOT complex, by RNAi-mediated knockdown in animals expressing an *ets-4* 3’UTR reporter. This reporter expresses, under the control of *ets-4* 3’UTR, histone H2B tagged with green fluorescence protein GFP (H2B::GFP), enabling the quantification of reporter expression by fluorescence microscopy. We found that NOT-1 and CCR-4 appeared dispensable for the *ets-4* silencing (Fig. 2D). Finally, we also RNAi-depleted PAN-2 and -3, which promote deadenylation independent of CCR4-NOT but, like above, found no effect on *ets-4* silencing (Fig. 2D). Collectively, our observations suggest that the silencing by REGE-1 requires RLE-1/Roquin, which appears to function independently from de-adenylation. Intriguingly, the functional interdependence between REGE-1 and RLE-1 suggests a mechanism involving cooperation between the two proteins, reminiscent of the initial model postulating mRNA co-regulation by Regnase-1 and Roquin-1 (4).

**Figure 2.**
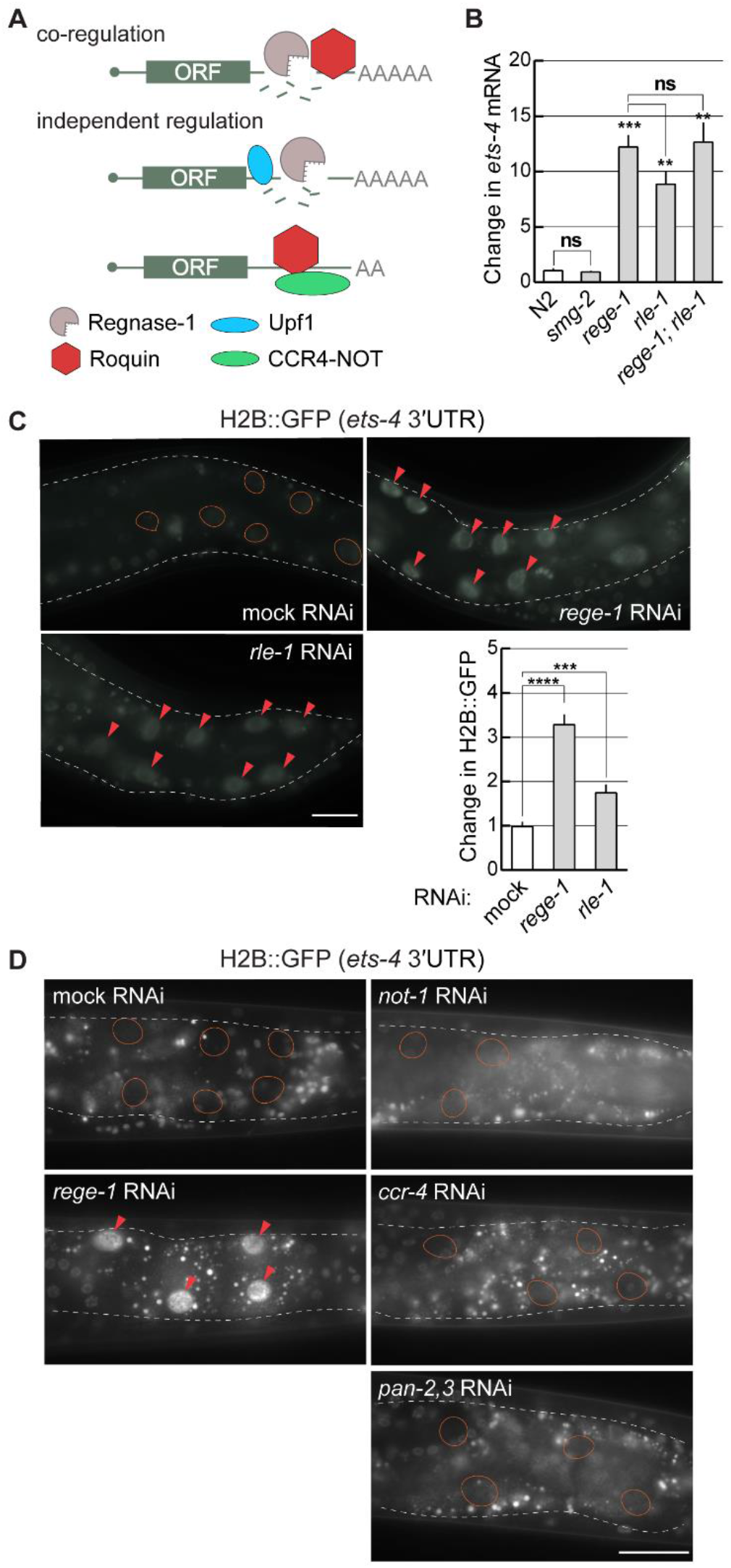
REGE-1 cooperates with RLE-1 in *ets-4* mRNA silencing. (A) Two models of mRNA silencing mediated by Regnase-1 and Roquin-1. Based on (4,5). The crucial difference between the two models is either cooperative (co-regulation), or independent functions of Regnase-1 and Roquin-1. In the co-regulation model, the mRNA-binding activity of Roquin-1 and the RNase activity of Regnase-1 are both essential for mRNA silencing. According to the independent model, Regnase-1 and Roquin-1 silence mRNAs independently of each other and they do so by different mechanisms: Regnase-1 regulates translationally active transcripts with the help of Upf1, while Roquin-1 controls translationally inactive mRNAs by recruiting the CCR4-NOT deadenylase complex. (B) RLE-1 is as important as REGE-1 for the *ets-4* mRNA silencing, while SMG-2/Upf1 does not seem to be involved. The level of *ets-4* mRNA was measured, by RT-qPCR, in animals of the indicated genotypes. Strains used: wt, *smg-2(rrr60), rege-1* (*rrr13*), *rle-1* (*rrr44*), and *rege-1* (*rrr13*); *rle-1* (*rrr44*) double mutant. The mRNA levels were normalized to the levels of *tbb-2* (tubulin) mRNA. Two-tailed P-values were calculated using an unpaired Student t-test. Bars represent the mean value from three independent biological replicates (N = 3). Error bars represent SEM; ** indicates P < 0.001, *** indicates P = 0.0001, “ns” = not significant. (C) RLE-1, like REGE-1, silences *ets-4* mRNA through its 3’ UTR. Upper left: partial view of live animals, with outlined intestines, expressing GFP::H2B reporter from a ubiquitous promoter (*dpy-30*), under the control of *ets-4* 3’ UTR. The animals were subjected to either mock, *rege-1* or *rle-1* RNAi, as indicated. Ovals mark the positions of gut nuclei not expressing the reporter GFP in control animals. Arrowheads indicate the gut nuclei in which, in wild type, the reporter is repressed. Scale bar: 20 µm. Lower right: The corresponding quantification of changes in the reporter GFP intensity. Between five and ten nuclei per animal, in at least five animals per condition, were analyzed, to quantify GFP intensities from thirty nuclei in total (N = 30). Two-tailed P-values were calculated using an unpaired Student t-test. Bars represent the mean value from GFP intensity. Error bars represent SEM. *** indicates P = 0.0001, **** indicates P < 0.0001. (D) Depletion of *C. elegans* deadenylases has no impact on *ets-4* silencing. Upper left: Partial view of live animals, expressing the GFP::H2B reporter from a ubiquitous promoter (*dpy-30*), under the control of *ets-4* 3’ UTR. Animals were subjected to either mock, *rege-1* (positive control), *ccr-4* or *not-1* (components of the CCR4-NOT complex), or *pan-2,3* (other deadenylases) RNAi, as indicated. Ovals mark the positions of gut nuclei not expressing the reporter GFP, arrowheads indicate the gut nuclei in which the reporter is repressed. Scale bar: 50 µm

### RLE-1 and REGE-1 have overlapping biological functions

We previously showed that REGE-1 facilitates the accumulation of body fat and cold resistance by lowering the levels of ETS-4 (Fig. 1A) (14,21). Because RLE-1, like REGE-1, is required for the silencing of *ets-4* mRNA, one could expect similar biological functions of both proteins. Indeed, we observed that *rle-1(-)* mutants, which, like *rege-1(-)*, overexpress ETS-4, have reduced levels of body fat and were cold-sensitive (Fig. 3A-B). Similar to *rege-1*, both of those *rle-1(-)* phenotypes were suppressed by the loss of ETS-4 (Fig. 3A-B). These results are consistent with the silencing of *ets-4* by both proteins. However, we noticed that the extend of cold survival appears to differ between the *rege-1(-)*; *ets-4(-)* and *rle-1(-)*; *ets-4(-)* double mutants, with the former surviving cold even better than wild type (21). It indicates that, in addition to *ets-4* co-regulation, RLE-1 and/or REGE-1 may play other roles that do not require their cooperation. The reported expression pattern of RLE-1 is consistent with this hypothesis (15). To confirm it, we obtained a strain expressing GFP-tagged RLE-1 from the endogenous locus. In agreement with the previous study, while the strongest expression of RLE-1::GFP was observed in the intestine, where REGE-1 is also expressed (14), we noticed additional, weaker but ubiquitous expression in other tissues (Fig. 3C). Thus, in addition to co-repressing *ets-4* with REGE-1, RLE-1 is expected to have additional functions.

**Figure 3.**
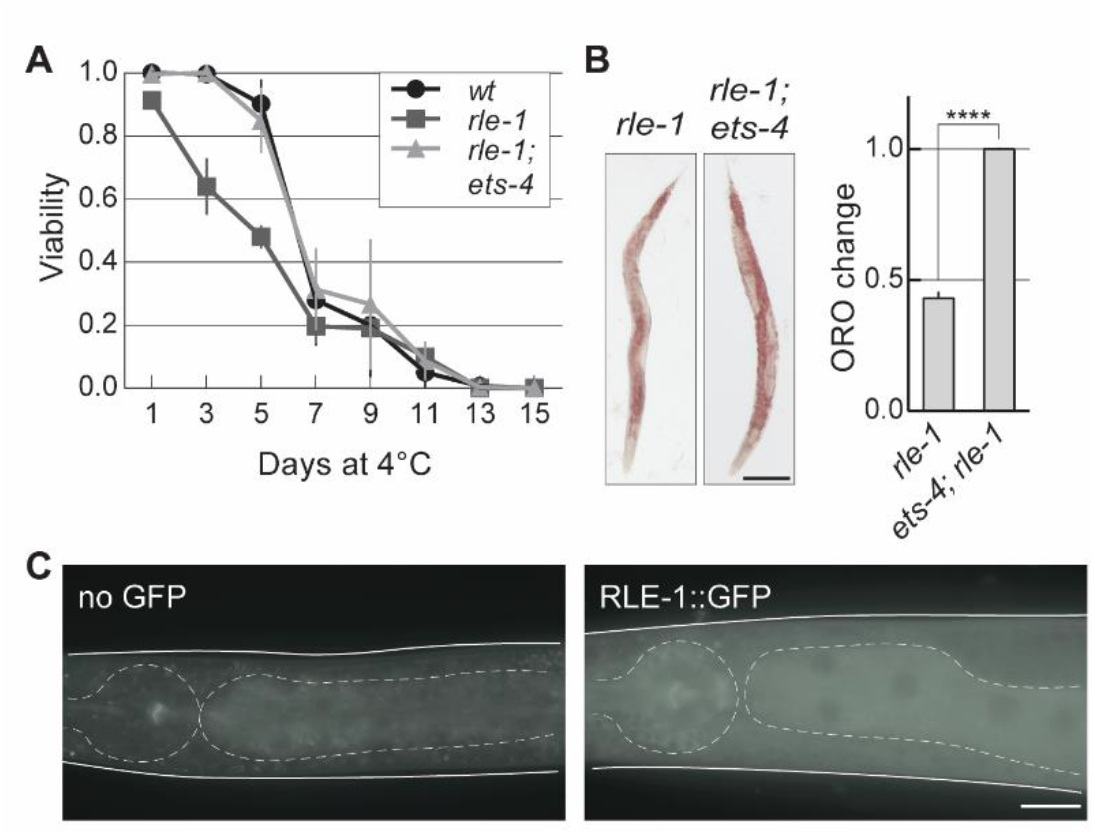
RLE-1 has similar biological roles to REGE-1. (A) RLE-1 is important for wild-type cold survival. Animals of the indicated genotypes were exposed to cold for the indicated time, and examined for viability one day after rewarming at 20°C. Tested were: wild type (wt; black), *rle-1(rrr44)* (dark grey), and *ets-4(rrr16); rle-1(rrr44)* double mutants (light grey). The experiment was performed in three biological replicates (N = 3); 100 animals were scored per time point. Bars represent the mean value from all replicates. Error bars represent SEM. (B) RLE-1 prevents ETS-4 dependent loss of body fat. Left: representative pictures of animals of the indicated genotypes with body fat stained with the lipophilic dye oil red O (ORO). Right: corresponding quantification of body fat. The experiment was performed in three biological replicates (N = 3); ten to fifteen animals were scored per replicate. Two-tailed P-values were calculated using an unpaired Student t-test. Bars represent the mean value from all biological replicates. Error bars represent SEM. *** indicates P = 0.0001. (C) RLE-1 is a ubiquitous protein enriched in the intestine. Partial view of representative live animals, which either did not (left) or did (right) express the endogenously tagged RLE-1::GFP protein. To reduce gut-specific autofluorescence, the animals carried the *glo-1(zu391)* mutation (26). As a control (no GFP) *glo-1* worms were used. The pharynx and intestines are outlined. Scale bar: 20 μm.

### The *ets-4* 3’UTR contains RNA stem-loops, ADE and RCE, which are required for the silencing

We showed previously that the degradation of *ets-4* mRNA by REGE-1 is mediated by a 115 nucleotide-long fragment of the *ets-4* 3’ UTR (the F1S fragment) (14). Roquin-1 is thought to bind RNA primarily via its ROQ domain (also present in RLE-1), which recognizes RNA SLs (8). Among these are the so-called ADE SLs, containing five nucleotides in the loop (11,12). By scanning the F1S sequence for putative regulatory motifs, we found an ADE SL, whose structural organization we have experimentally confirmed by SHAPE (Selective 2′-hydroxyl acylation analyzed by primer extension) (Fig. 4A, S2). The sequence in the apical loop of the *ets-4* ADE SL conforms to the ADE SL consensus recognized by the ROQ domain of Roquin-1 (12), and is almost identical to the ADE SL found in the *OX40* 3’UTR (Fig. 4B) (11). Intriguingly, the *ets-4* ADE SL is found next to another SL, which we called the RCE (<u>R</u>EGE-1 <u>c</u>leavage <u>e</u>lement), containing the REGE-1 cleavage site (Fig. 4A) (14). Thus, RLE-1 could repress *ets-4* via the association with the ADE SL within the F1S fragment of *ets-4* 3’UTR.

**Figure 4.**
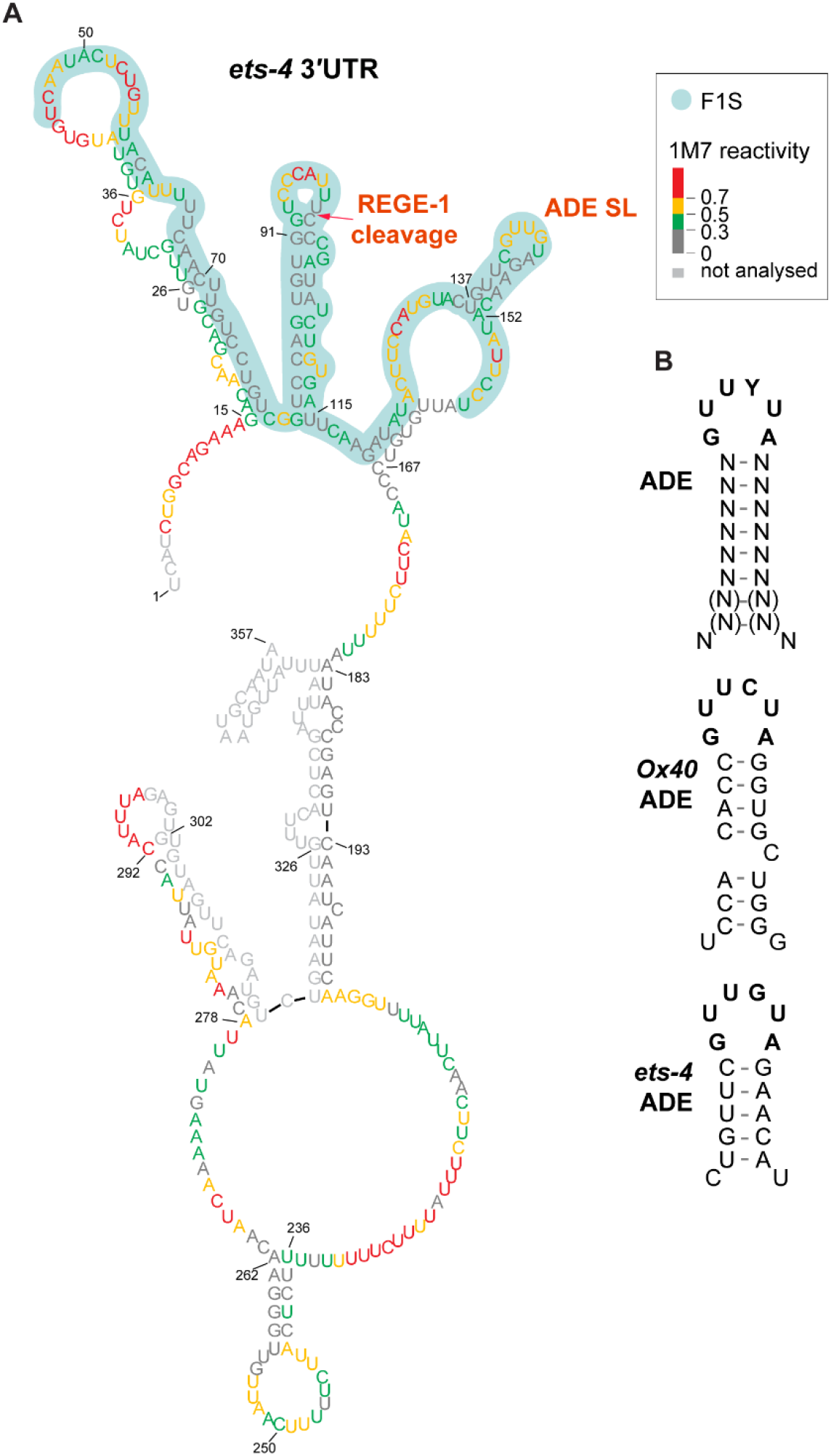
The F1S fragment of *ets-4* 3’ UTR contains ADE SL, potentially mediating the repression by RLE-1. (A) The F1S fragment of *ets-4* 3’ UTR contains an ADE SL. Shown is the *ets-4* 3’ UTR structure obtained by SHAPE *in vitro* probing. The RNA reactivity patterns (labelled as 1M7 reactivity) were obtained for the isolated F1S sequence or in the context of the whole *ets-4* 3’ UTR. The F1S encompasses a 115 nt fragment, between the residues U45 and U159 (highlighted in light green). The red arrow indicates the REGE-1 cleavage site, based on (14). The consensus motif in the ADE stem-loop (Alternative Decay Element) is indicated as ADE SL. (B) The ADE SL from *ets-4* 3’ UTR is almost identical to the motif recognized by Roquin-1. Top: mammalian consensus for ADE stem-loop structural motif recognized by mammalian Roquin-1; based on (12), middle: ADE type stem-loop from mammalian mRNA *OX40*, bottom: ADE-like stem-loop from worm *ets-4* mRNA.

Thus far, we showed that the F1S fragment of *ets-4* 3’UTR is sufficient for the repression by REGE-1 (14). Therefore we tested whether that repression, similar to the full-length 3’UTR, also depends on RLE-1. Indeed, we found that repression of a GFP reporter, mediated by the F1S fragment of *ets-4* 3’UTR inserted into otherwise unregulated *unc-54* 3’UTR, required both REGE-1 and RLE-1 (Fig. 5A). These results suggest that the F1S fragment contains all information required for the silencing by both proteins (Fig. 5B). We then asked whether the ADE and RCE SLs are important for silencing. To test it, we created versions of the F1S GFP reporter, carrying deletions of either the ADE (F1S ΔADE) or RCE (F1S ΔRCE) SL (Fig. 5C). Monitoring the expression of these reporters, we found that the deletion of each SL resulted in the reporter de-repression (Fig. 5C). Collectively, our data suggest a model where the ADE element contributes to the silencing by recruiting RLE-1, and the RCE element by facilitating, in some way, the cleavage by REGE-1.

**Figure 5.**
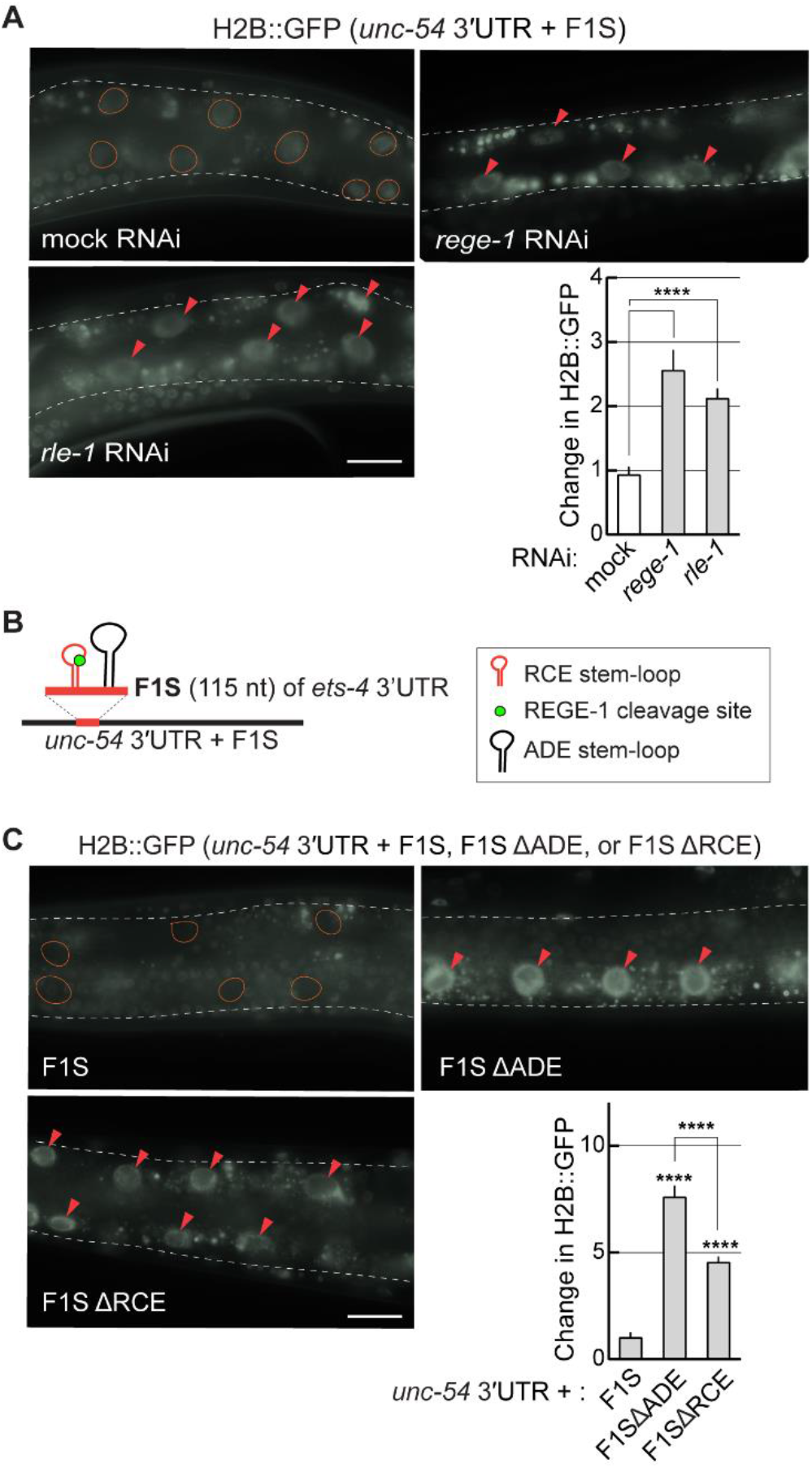
The F1S fragment is sufficient for REGE-1 and RLE-1 mediated mRNA regulation. (A) The repression mediated by the F1S fragment requires both REGE-1 and RLE-1. Upper left: partial view of live, wild-type animals, with outlined intestines, expressing the GFP::H2B reporter under the control of a modified *unc-54* 3’ UTR (containing the F1S fragment of *ets-4*, indicated as the *unc-54* 3’ UTR + F1S). The animals were subjected to either mock, *rege-1*, or *rle-1* RNAi, as indicated. Ovals mark positions of gut nuclei in which, in wild type, the reporter is repressed. Arrowheads indicate the gut nuclei expressing the reporter GFP. Scale bar: 20 µm. Bottom right: the corresponding quantification of changes in the GFP intensity. Between five and ten nuclei per animal, in at least five animals per condition, were analyzed, to get GFP intensities from thirty five nuclei in total (N = 35). Two-tailed P-values were calculated using an unpaired Student t-test. Bars represent the mean value from GFP intensity. Error bars represent SEM. *** indicates P = 0.0001, **** indicates P < 0.0001. (B) Schematic representation of a hybrid 3’UTR, controlling expression of a GFP reporter. The F1S fragment from the *ets-4* 3’ UTR, which mediates REGE-1 induced RNA degradation, was ‘‘transplanted’’ into an otherwise unregulated 3’ UTR, resulting in a “hybrid” 3’UTR (*unc-54* 3’ UTR + F1S). The F1S was inserted between bps 164 and 165 of the *unc-54* 3’ UTR; the F1S contains the REGE-1 cleavage site (RCE stem-loop) and the ADE SL (ADE stem-loop). (C) The deletions of either ADE or RCE result in the reporter de-repression. Upper left: partial view of live, wild-type animals, with outlined intestines, expressing the GFP::H2B reporter under the control of the modified *unc-54* 3’ UTR, containing the F1S fragment of *ets-4* 3’ UTR or its versions, with deleted ADE SL (ΔADE) or the REGE-1 cleavage site (ΔRCE). Ovals demarcate the gut nuclei in which, in wild type, the reporter is repressed. Arrowheads indicate the gut nuclei expressing the reporter GFP. Scale bar: 20 µm. Lower right: the corresponding quantification of changes in the GFP intensity. Between five and ten nuclei per animal, in at least five animals per condition, were analyzed to quantify thirty-five nuclei in total (N = 35). Error bars represent SEM. An unpaired two-tailed t-test was used to calculate the P-value. The *unc-54* 3’ UTR + F1S (F1S for short) was used as a reference. Lines above the bars indicate values that were used for comparison; **** indicates P > 0.0001.

### RLE-1 binds the ADE-like SL via its ROQ domain

Since the ROQ domain is conserved between RLE-1 and Roquin-1 (Fig. 6A), and is known to bind ADE SLs (11,12), we wondered whether this domain also mediates the binding of RLE to the F1S ADE. To examine that, we created a strain expressing, from the endogenous locus, a mutant form of RLE-1, carrying amino acid substitutions (Y245A, K254A, R255A) in the ROQ domain (RLE-1 ROQmut), as the corresponding substitutions in Roquin-1 (Y250A, K259A, and R260A) abolish the RNA binding (11,25). We found that, in the strain expressing the RLE-1 ROQmut, neither the endogenous *ets-4* mRNA nor the F1S reporter was repressed (Fig. 6B-C). To examine the interaction between the RLE-1 ROQ domain and the *ets-4* ADE, we initially attempted to immunoprecipitate RLE-1 but, possibly due to the low levels of RLE-1, were unable to detect the protein. We thus synthesized RLE-1 (or its ROQmut variant) in reticulocyte lysates and, using a biotinylated F1S fragment (or its mutated versions) as a bait, performed RNA pulldowns, monitoring the interaction with RLE-1 variants by western blotting. With this approach, we observed the interaction between RLE-1 and the F1S RNA. Importantly, this interaction was lost when the F1S ADE (but not RCE) was deleted, or when we tested the RLE-1 ROQmut protein (Fig. 6D). Using the same approach, we attempted to examine the interaction between the F1S RNA and REGE-1 but, at least *in vitro*, REGE-1 associates with RNA non-specifically, independently of RLE-1 (data not shown). However, we have shown in the past that, *in vivo*, the RNase-dead REGE-1 associates with the *ets-4* mRNA specifically (14). Thus, we performed immunoprecipitations between RNase-dead REGE-1 and the endogenous *ets-4* mRNA, and tested whether their interaction depends on RLE-1. Importantly, we observed that REGE-1 was able to associate with *ets-4* mRNA both in the presence or absence of RLE-1 (Fig. 6E). Taken together, our data suggest that the binding of the two proteins to *ets-4* is independent of each other.

**Figure 6.**
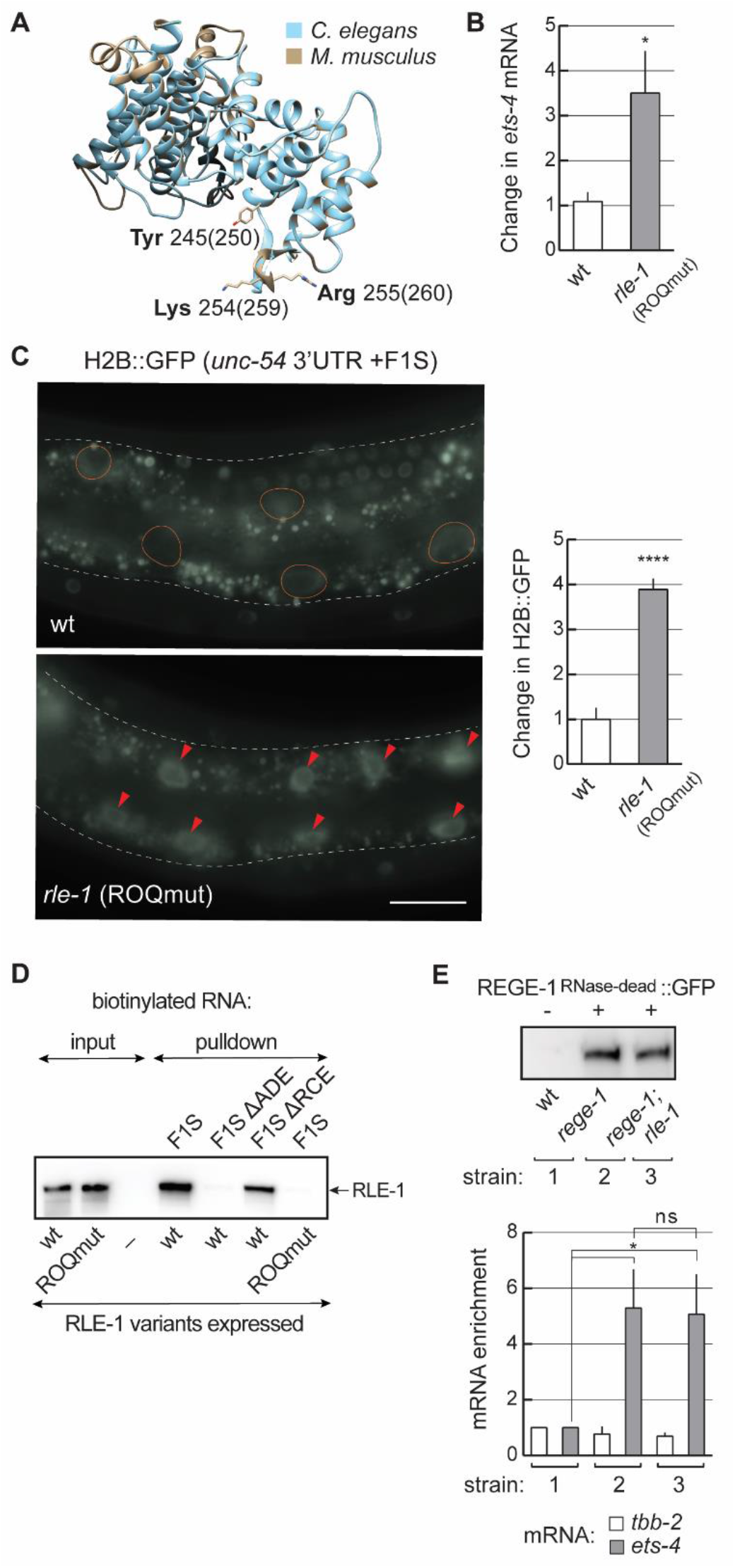
The ROQ domain of RLE-1 is essential for *ets-4* silencing. (A) The ROQ domain is conserved between RLE-1 and Roquin-1. A homology model of the *C. elegans* RLE-1 ROQ domain. The RLE-1 ROQ domain tertiary structure was predicted based on conservation with the ROQ domain of mouse Roquin-1 (25), using Phyre2 software. The RLE-1 ROQ domain is in light blue and the Roquin-1 ROQ domain is in grey. Amino acids mutated in the ROQmut (Tyr245, Lys254, Arg255) are indicated. (B) Mutations in the ROQ domain (Y245A, K254A, R255A) result in *ets-4* mRNA de-repression. The levels of endogenous *ets-4* mRNA were measured, by RT-qPCR, in animals of the indicated genotypes. Strains used: wt, *rle-1* (*syb517)* - expressing the RLE-1 ROQmut. Bars represent the mean from five biological replicates (N = 5), error bars represent SEM. Two-tailed P-values were calculated using unpaired Student t-test, wt was used as a reference.* indicates P < 0.05. (C) Mutations in the RLE-1 ROQ domain cause de-repression of the F1S reporter. Left: partial view of live animals, either wild-type or expressing RLE-1 with substitutions in the ROQ domain (ROQmut: Y245A, K254A, R255A) (as in B), and expressing the GFP::H2B reporter under the control of a modified *unc-54* 3’ UTR (*unc-54* + F1S). The intestines were outlined, ovals demarcate the gut nuclei in which, in wild type, the reporter is repressed, arrowheads indicate the gut nuclei expressing the reporter GFP. Scale bar: 20 µm. Right: the corresponding quantification of changes in the GFP intensity. Between five and ten nuclei per animal, in at least five animals per condition, were analyzed, to quantify GFP intensities from thirty-five nuclei in total (N = 35). Two-tailed P-values were calculated using an unpaired Student t-test, wt was used as a reference. Bars represent the mean value from GFP intensity. Error bars represent SEM.**** indicates P < 0.0001. (D) The ADE stem-loop is crucial for the RNA binding by RLE-1, and mutations in the ROQ domain abolish RLE-1 interaction with the F1S RNA. *In vitro* synthesized, biotinylated RNAs (F1S, F1S ΔADE or F1S ΔRCE) were pulled down after incubation with *in vitro*-translated wild-type RLE-1, or the RLE-1 mutant (ROQmut), carrying amino acid substitutions (Y245A, K254A, R255A) in the ROQ domain. Proteins associated with biotinylated RNAs were examined with a western blot. The experiment was performed three times (N = 3). (E) REGE-1 associates with the *ets-4* mRNA independently of RLE-1. Animals of the indicated genotypes were lysed and the extracts were subjected to immunoprecipitation (IP), using anti-GFP antibodies. Top: western blot of immunoprecipitates from wt or mutant animals; *rege-1(rrr13)* or *rege-1(rrr13); rle-1(rrr44)* mutants that express the RNase-dead REGE-1::GFP, detected with the REGE-1 antibody (14), was successfully immunoprecipitated from the lysates with similar efficiency. Below: the corresponding quantification of the indicated mRNAs, by qRT-PCR, which co-precipitated with the RNase-dead REGE-1. Note that the RNase-dead REGE-1::GFP is associated with the *ets-4* mRNA independently of RLE-1. The mRNA levels were normalized to the levels of *tbb-2* (tubulin) mRNA. Bars represent the mean from four biological replicates (N = 4), error bars represent SEM. Two-tailed P-values were calculated using unpaired Student t-test; lines above the bars indicate values that were compared; “ns” = not significant, * indicates P < 0.05.

### Like REGE-1 and RLE-1, the ADE and RCE SLs are equally important for the *ets-4* decay

Our data suggest that RLE-1 associates with *ets-4* via the ADE and REGE-1 cleaves the RNA within the RCE SL. To test the relative contribution of these RNA elements, we examined to what degree the loss of ADE or RCE impacts the silencing. To do that, we compared the expression of F1S ΔADE and F1S ΔRCE reporters in the presence or absence of REGE-1 or RLE-1. Since RLE-1 appears to associate with *ets-4* 3’UTR via the ADE, we found, not surprisingly, that the loss of RLE-1 did not additionally increase the expression of the F1S ΔADE reporter (Fig. 7A). Interestingly, however, this reporter was also insensitive to the loss of REGE-1 (Fig. 7A). Thus, even though REGE-1 binds *ets-4* 3’UTR independently of RLE-1, the interaction between RLE-1 and ADE is essential for the silencing by REGE-1. We then examined the F1S ΔRCE reporter and found that, like the deletion of ADE, the deletion of RCE completely abolished the regulation by REGE-1 and RLE-1, as the loss of either protein did not impact the expression of the reporter (Fig. 7B).

**Figure 7.**
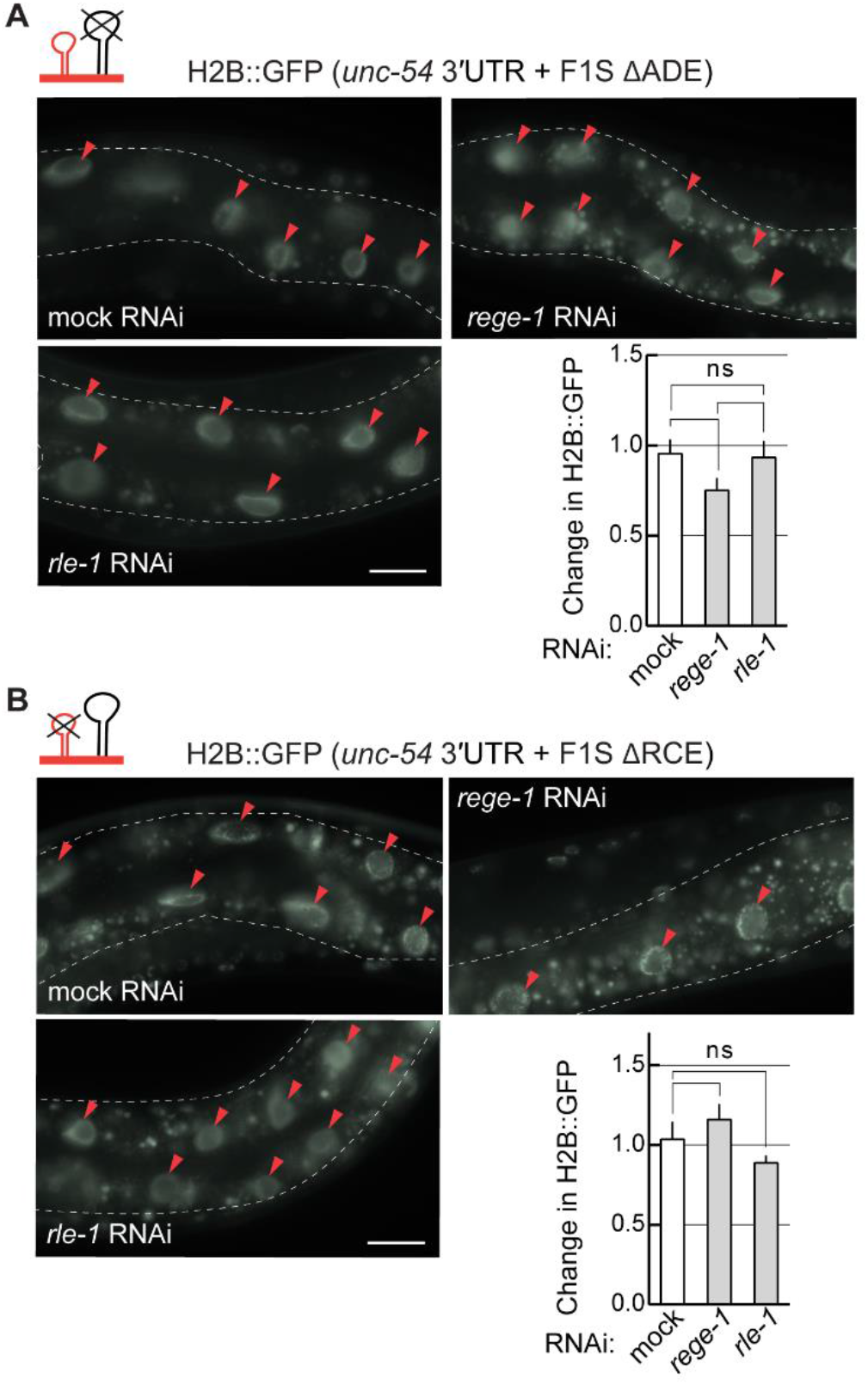
The ADE and the RCE stem-loops are essential for *ets-4* mRNA regulation. (A) The reporter lacking ADE is no longer regulated by REGE-1 or RLE-1. Top: the mutated F1S version was ‘‘transplanted’’ into an otherwise unregulated 3’ UTR (*unc-54* + F1S ΔADE; the fragment was inserted between bps 164 and 165 of the *unc-54* 3’ UTR). Upper left: Partial view of live, wild-type animals, with outlined intestines, expressing the GFP::H2B reporter under the control of a modified *unc-54* 3’ UTR (containing the F1S fragment with deleted ADE (ΔADE). The animals were subjected to either mock, *rege-1*, or *rle-1* RNAi, as indicated. Ovals mark gut nuclei in which, in wild type, the reporter is repressed. Arrowheads indicate the gut nuclei expressing the reporter GFP. Scale bar: 20 µm. Bottom right: the corresponding quantification of changes in the GFP intensity. Between five and ten nuclei per animal, in at least five animals per condition, were analyzed, to quantify GFP intensities from thirty-five nuclei in total (N = 35). Two-tailed P-values were calculated using an unpaired Student t-test. Bars represent the mean value from GFP intensity. Error bars represent SEM. “ns” indicates not significant. The RCE is essential for REGE-1 and RLE-1 mediated silencing of the F1S reporter. Same as in B, except that the REGE-1 cleavage element was deleted (ΔRCE). Between five and ten nuclei per animal, in at least five animals per condition, were analyzed, to get GFP intensities from thirty nuclei in total (N = 30). Two-tailed P-values were calculated using an unpaired Student t-test. Bars represent the mean value from GFP intensity. Error bars represent SEM. “ns” indicates not significant.

The above experiments showed the contribution of ADE and RCE to *ets-4* silencing. To directly test if RLE-1 and ADE are as important for the *ets-4* degradation as are REGE-1 and RCE, we designed primers to indirectly monitor (by qRT-PCR) the RNA cleavage (Fig. 8A-B). We found the levels of *ets-4* mRNA fragments spanning the REGE-1 cleavage site were as abundant in *rege-*1 mutants as in *rle-1* or *rle-1; rege-1* mutants (Fig. 8A). These results suggest that *ets-4* mRNA does not undergo REGE-1 mediated cleavage in the absence of RLE-1. We additionally performed the same analysis on the F1S reporters and found that the deletion of RCE had the same effect on the levels of *ets-4* mRNA fragments (spanning the REGE-1 cleavage site) as the deletion of ADE (Fig. 8B). Taken together, our findings support a model, where REGE-1 and RLE-1 (R2) bind the *ets-4* 3’UTR independently but collaborate to create a local microenvironment promoting, possibly with the help of additional factors, the RNA cleavage by REGE-1 (Fig. 8C).

**Figure 8.**
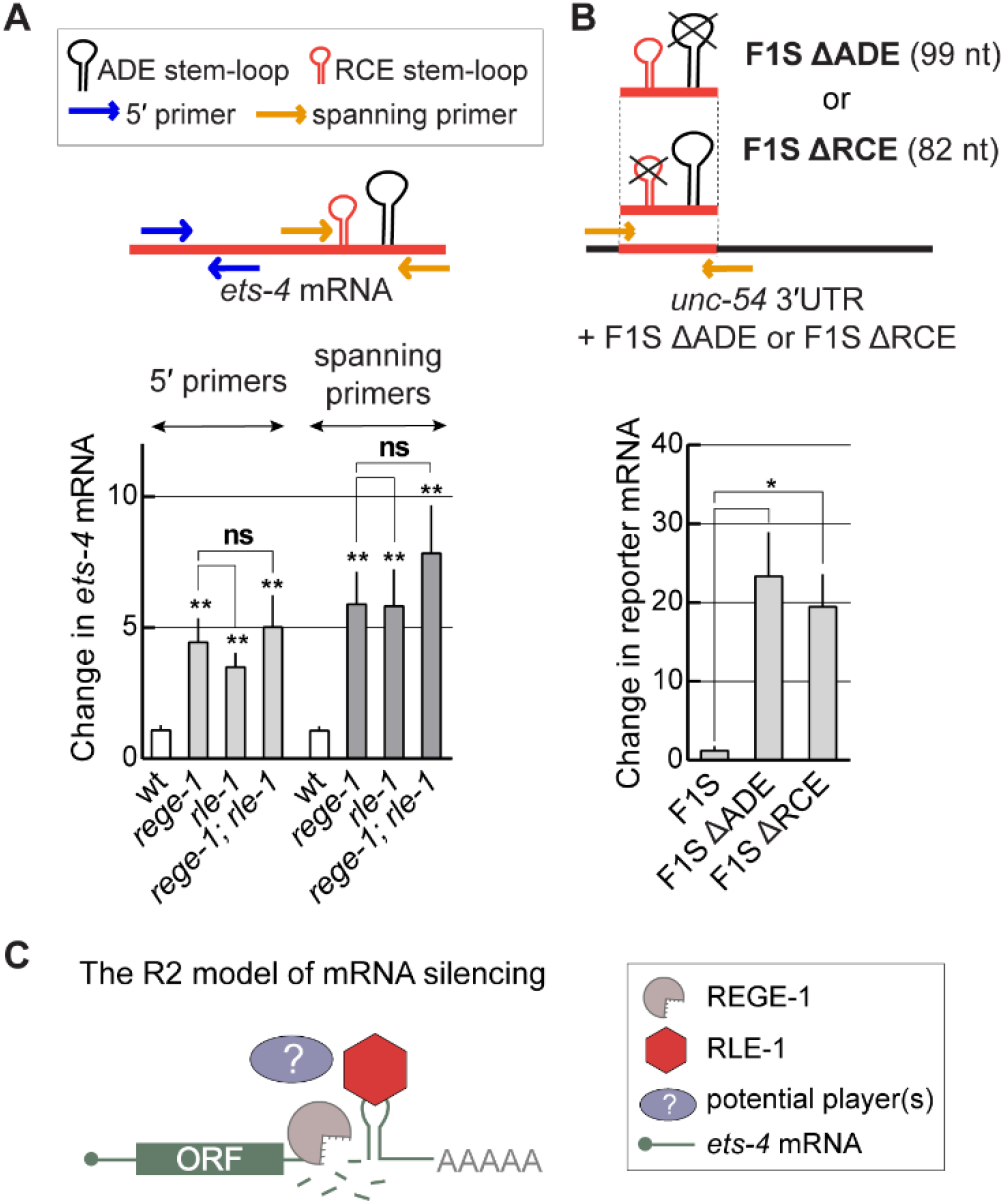
RLE-1 is crucial for REGE-1 mediated RNA cleavage. (A) *ets-4* mRNA fragments spanning the REGE-1 cleavage site were as abundant in the absence of REGE-1 as in the absence of RLE-1. Top: schematic representation of the *ets-4* 3’UTR and RT-qPCR primers designed to amplify fragments upstream (blue arrows) or spanning the REGE-1 cleavage site (orange arrows). Bottom: the levels of *ets-4* mRNA were measured, by RT-qPCR, in animals of the indicated genotypes. Strains used: wt, *rege-1* (*rrr13*), *rle-1* (*rrr44*), and *rege-1* (*rrr13*); *rle-1* (*rrr44*) double mutant. The mRNA levels were normalized to the levels of *tbb-2* (tubulin) mRNA. Bars represent the mean from six biological replicates (N = 6), error bars represent SEM. Two-tailed P-values were calculated using unpaired Student t-test compared to wt. Lines above the bars indicate values that were compared. “ns” = not significant, ** indicates P < 0.001. (B) The F1S reporter lacking the ADE or the RCE SL is no longer degraded. Top: schematic representation of *unc-54* 3’UTR + F1S ΔADE or F1S ΔRCE, and the RT-qPCR primers designed to amplify fragment spanning the REGE-1 cleavage site (orange arrows). The levels of F1S, F1S ΔADE, and F1S ΔRCE reporter mRNAs were measured, by RT-qPCR. Bars represent the mean from three biological replicates (N = 3), error bars represent SEM. Two-tailed P-values were calculated using unpaired Student t-test, lines above the bars indicate values that were compared. Error bars represent SEM, * indicates P < 0.05. (C) The proposed model of *ets-4* mRNA silencing, mediated by REGE-1 and RLE-1 (R2). Each protein binds the *ets-4* mRNA independently of each other, but the binding of both proteins is crucial for mRNA decay. According to this model, which we dubbed the “R2 model of mRNA silencing”, the ADE-recruited RLE-1 is essential for mRNA-associated REGE-1 to cleave the mRNA, leading to its destruction. Whether REGE-1 binds to mRNA alone, or in association with other proteins, remains unclear. Also, it remains to be determined how exactly RLE-1 promotes the RNase activity of REGE-1.

## DISCUSSION

We describe here the mechanism of REGE-1 mediated *ets-4* mRNA decay dependent on the RBP RLE-1. Here, similar to Roquin-1, RLE-1 binds RNA via the interaction between its ROQ domain and the ADE SL. In mammals, when functioning independently from Regnase-1, Roquin silences its mRNA targets by recruiting the CCR4-NOT deadenylase complex (5,8). Human Roquin-1, Roquin-2, and the fruit fly Roquin interact with the CCR4-NOT complex via their unstructured C-terminal regions, despite the lack of sequence conservation (4,24). The recruitment of CCR4-NOT by the fruit fly Roquin is mediated by CAF binding motif (CBM). However, Roquin proteins from other insects, nematodes, and vertebrates do not carry a recognizable CBM (24). Thus, the recruitment of CCR4-NOT may not be a universal feature of all Roquin-related proteins, possibly explaining why the *ets-4* repression by RLE-1 does not involve the CCR4-NOT complex. Instead, we found that the RLE mediated silencing requires REGE-1. While REGE-1 induces RNA cleavage in the close vicinity of the ADE-associated RLE-1, the recruitment of REGE-1 to the cleavage element, RCE, remains a conundrum. We failed, thus far, to detect a direct interaction between REGE-1 and RLE-1, arguing against RLE-1 mediated recruitment of REGE-1 to *ets-4* mRNA. The RLE-1-independent recruitment of REGE-1 is further supported by the observation that REGE-1 still associates with *ets-4* in the absence of RLE-1. Then, where does the mRNA specificity of REGE-1 come from, and why the RNA cleavage by REGE-1 requires RLE-1? Regnase-1 targets overlap with Roquin targets and both RBPs require similar SL structures to associate with RNAs (5,8,12). However, whether Regnase-1 association with RNA is direct, or depends on other factors, remains unclear. *In vitro*, both Regnase-1 and REGE-1 display no specificity towards RNAs that, *in vivo*, are specifically regulated (10) (our data not shown). Thus, we speculate that the specificity of REGE-1 towards *ets-4* may depend on another protein, recruiting REGE-1 to the 3’UTR. As for the possible functional link between RLE-1 and REGE-1, RLE-1 might impact a local RNA/protein conformation in a way that promotes the cleavage by REGE-1, potentially by enabling the interaction between the REGE-1 active site with the *ets-4* RCE. There are, however, other possible scenarios and future work, likely involving *in vitro* reconstitution of the RNA cleavage, may be necessary to understand the functional relationship between the contributing factors.

Intriguingly, the functional interdependence, similar to that between REGE-1 and RLE-1, was originally reported between Regnase-1 and Roquin-1 (4). Because a chimeric protein, harboring the ROQ-domain of Roquin-1 fused to Regnase-1, was sufficient to regulate mRNA reporter in Roquin-1 or Regnase-1 deficient mouse embryonic fibroblasts (MEF) cells (4), Roquin-1 was proposed to recruit Regnase-1 to mRNA. In these studies, the repression depended on the RNA-binding domain of Roquin and the RNase activity of Regnase-1 (4). However, a physical interaction between the wild-type proteins has not been demonstrated. Nonetheless, similar to our observations on *ets-4*, the repression of at least some mammalian mRNAs, like *Nfkbid, Nfkbiz, cRel*, and *Irf4*, requires both Regnase-1 and Roquin (4). Also, reporter studies showed that neither Roquin nor Regnase-1, which was expressed without the other protein in Roquin and Regnase-1 deficient MEF cells, was sufficient to repress reporters controlled by *Ctla4, Icos*, and *cRel* 3′ UTRs (4). Seemingly, these results contradict other studies that demonstrated the independent functions of Regnase-1 and Roquin (5,6). However, that discrepancy could be explained by the existence of different mechanisms, where the choice of a mechanism may depend on a cellular environment and/or mRNA target. While the data supporting the co-regulation by Regnase-1 and Roquin comes from MEFs and mouse T cells (4), the independent functions of the two proteins are supported by studies on other cell types (MEF cells, macrophages, HeLa cells, HEK cells, Jurkat cells, and NIH/3T3 cells) (5,6). Thus, assuming that Regnase-1 and Roquin can co-regulate some targets, the interdependence could underlie an ancient silencing mechanism operating from nematodes to humans. According to this hypothesis, in mammals, Regnase-1 and Roquin acquired additional, independent functions. However, the opposite is also possible, with the two proteins evolving to co-regulate some targets under certain circumstances. Although thus far, the overexpression of ETS-4 appears to explain all phenotypes associated with the loss of REGE-1 activity, we cannot exclude the existence of other REGE-1 targets that might be regulated independently of RLE-1. Conversely, RLE-1 is likely to regulate other mRNAs independently of REGE-1, as it is expressed more broadly than REGE-1. Whether/how RLE-1 controls mRNAs in tissues devoid of REGE-1 is an interesting question for future studies.

Irrespective of the molecular mechanisms employed by REGE-1/Regnase-1 and RLE-1/Roquin, it is intriguing that these proteins target mRNAs encoding proteins involved in the immune response. This is despite major differences in molecular mechanisms underlying nematode versus mammalian immunity. Moreover, REGE-1 and RLE-1 appear to regulate immune response indirectly, by silencing a transcription factor regulating the expression of many defence genes (14). By contrast, Regnase-1 and Roquin target mRNAs encoding factors, like the cytokines IL-2, IL-6, and TNF, or the T-cell co-stimulatory ICOS and OX40, which directly regulate the immune response (3). Thus, either the conserved biological roles of REGE-1/Regnase-1 and RLE-1/Roquin in the immune defence are accidental or, more interestingly, could reflect an ancient function of these proteins in a primordial immunity of ancestral organisms.

## ACKNOWLEDGMENTS

We are grateful to prof. Osamu Takeuchi (KU) for sharing mRegnase-1 and mRegnase-1 D141N expression vectors, and reporter plasmid for *IL6* expression. We thank Katarzyna Purzycka (IBCh PAS) for facilitating the collaboration with KPW and JG, Weronika Wendlandt-Stanek (IBCh PAS) for the modeling of ROQ domains, Aneta Dyczkowska (IBCh PAS) for the ORO staining, Bogna Juskowiak (IBCh PAS) for general technical help, and Jedrzej Malecki (UiO) for comments on the manuscript.

