## Supplemental information for "An mRNA silencing mechanism reliant on the cooperation between REGE-1/Regnase-1 and RLE-1/Roquin-1"

### SUPPLEMENTARY FIGURES

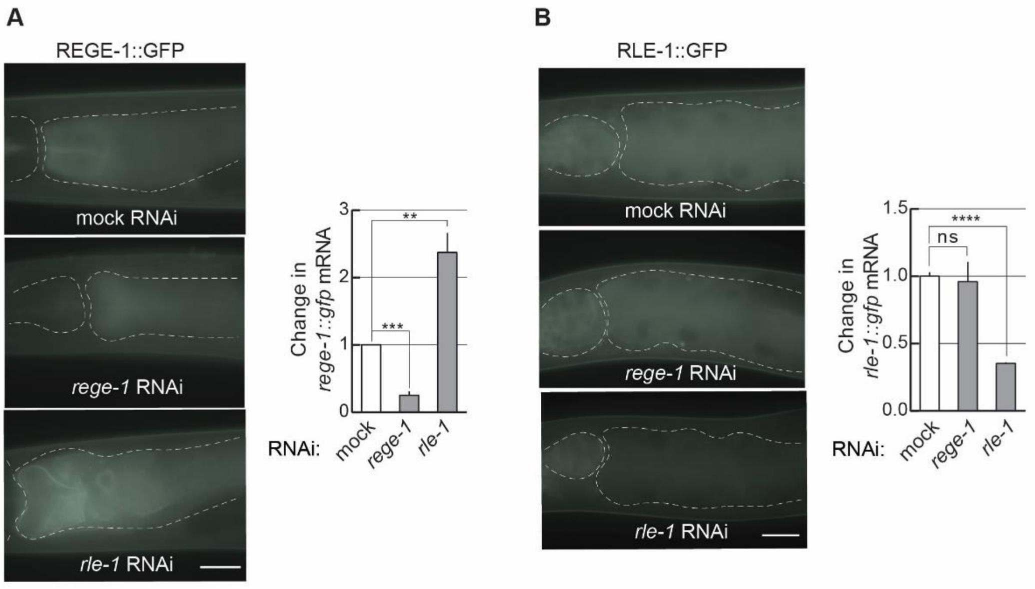


**Figure S1. REGE-1 expression increases in the absence of RLE-1, while the expression of RLE-1 is not affected by the loss of REGE-1.**

(A) Depletion of RLE-1 leads to REGE-1 upregulation. Left: partial view of representative live animals expressing rescuing REGE-1::GFP, subjected to either mock, *rege-1* or *rle-1* RNAi, as indicated. To reduce gut-specific autofluorescence, the animals carried the *glo-1*(zu391) mutation (1). The pharynx and intestines are outlined. Scale bar: 20 μm. Right: The corresponding quantification of changes in *rege-1::gfp* mRNA measured by qRT-PCR. Bars represent the mean value from three independent biological replicates (N = 3). Error bars represent SEM; Two-tailed P-values were calculated using unpaired Student t-test using mock RNAi as a reference. ** indicates $P < 0.001$, *** indicates $P = 0.0001$.

(B) Depletion of REGE-1 does not impact the levels of RLE-1. The same as in A, except that animals expressed endogenously GFP-tagged RLE-1 protein, and changes in *rle-1::gfp* mRNA were measured. Bars represent the mean value from three independent biological replicates (N = 3). Error bars represent SEM; Two-tailed P-values were calculated using unpaired Student t-test using mock RNAi as a reference. **** indicates $P < 0.0001$.

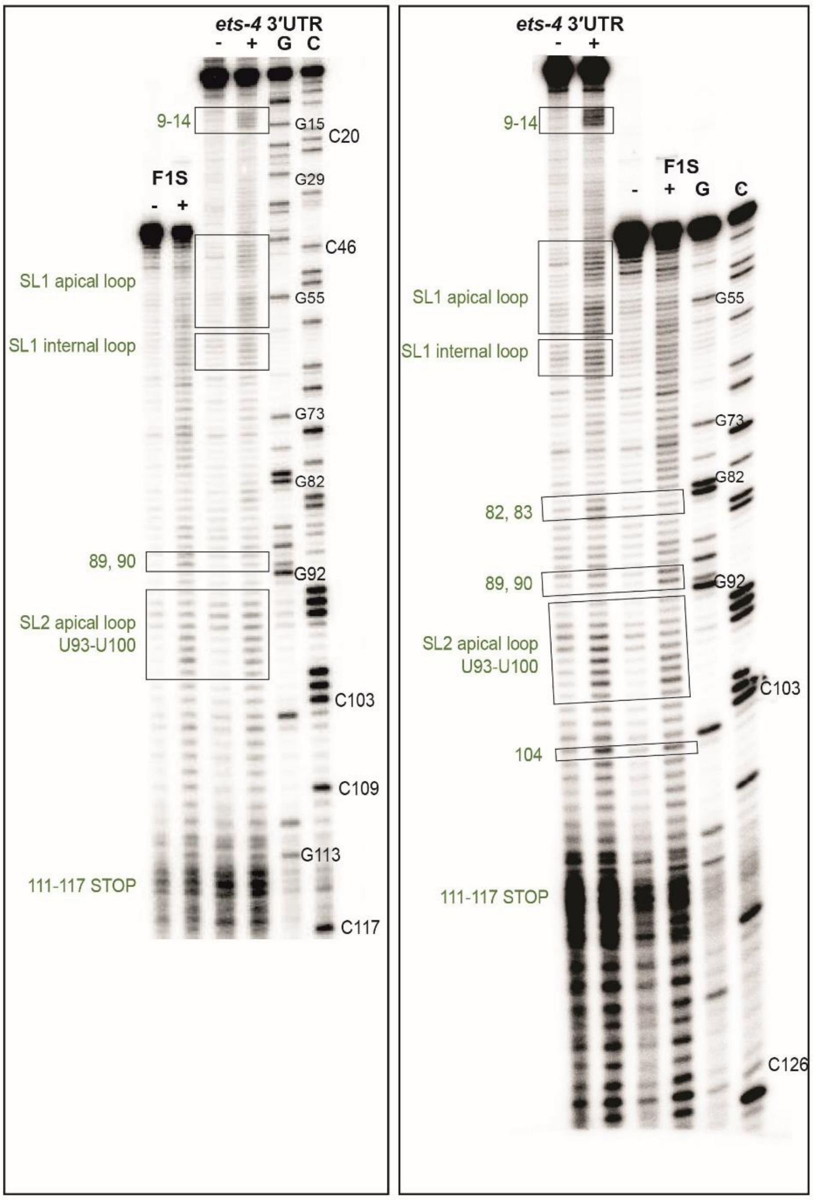


**Figure S2. Gel electrophoresis fractionation of products resulting from SHAPE structure probing experiments of F1S and *ets-4* 3'UTR.**

The F1S fragment of *ets-4* 3'UTR contains an ADE stem-loop. Two independent, representative gels from SHAPE structure probing are presented. Both RNAs were run concurrently to allow direct comparison. Lane (−) represents the control sample with untreated RNA; lane (+) 1M7 modification, C and G are sequencing lanes.

### SUPPLEMENTARY TABLES

**Table S1. The *C. elegans* strains used in this work.**

| **Genotype** | **CGC**/RAF |
|---|---|
| N2 Bristol | **N2** |
| *smg-2 (rrr60) I. [Q265*]* | 5097 |
| *rege-1(rrr13) I.* | 1657 |
| *rle-1 (rrr44) III.* | 2055 |
| *rege-1(rrr13) I; rle-1 (rrr44) III.* | 2102 |
| *rrrSi412 [Pdpy-30::gfp::h2b::ets-4 3'UTR; unc-119 (+)] II.;unc-119(ed3) III.* | 1740 |
| *rrrSi428[Pdpy-30::gfp::h2b::unc-54 3'UTR with transplanted (F1S) of ets-4 3`UTR; unc-119(+)] II; unc-119(ed3) III.* | 1844 |
| *rrrSi500[Pdpy-30::gfp::h2b::unc-54 3'UTR with transplanted mutated (F1SΔADE) of ets-4 3`UTR; unc-119(+)] II; unc-119(ed3) III.* | 5076 |
| sybSi111*[Pdpy-30::gfp::h2b::unc-54 3'UTR with transplanted mutated (F1SΔRCE) of ets-4 3`UTR; unc-119(+)] II; unc-119(ed3) III.* | 5113 |
| *glo-1(zu391) X.* | **JJ1271** |
| *glo-1(zu391) X.; rrrSi411 [Prege-1::rege-1(cDNA)::gfp::rege-1 3'UTR; unc-119(+)] II.* | 1761 |
| *glo-1(zu391) X.; rle-1 (syb1279) III* | 5120 |
| *rrrSi431 [Prege-1::rege-1(cDNA [D231N, D313A, D314A, D332A])::gfp::rege-1 3'UTR; unc-119(+)] II.;unc-119(ed3) III.; rege-1(rrr13) I.* | 5046 |
| *rrrSi431 [Prege-1::rege-1(cDNA [D231N, D313A, D314A, D332A])::gfp::rege-1 3'UTR; unc-119(+)] II.;unc-119(ed3) III.; rege-1(rrr13) I.; rle-1 (rrr44) III.* | 5047 |
| *rle-1 (syb1279) III; rege-1(rrr13) I.* | 5077 |
| *rle-1(syb517) III.* | 5003 |
| *ets-4(rrr16) X.* | 1758 |

“CGC” indicates strain numbers deposited in the Caenorhabditis Genetics Center; “RAF” strain numbers in the Ciosk lab collection.

**Table S2. DNA oligonucleotides used in this study.**

| Purpose | Name | Sequence |
|---|---|---|
| qPCR primers | qPCR_ets-4 F up | CTGAGAACCCGAATCATCCA |
| | qPCR_ets-4 R up | TCATTCATGTCTTGACTGCTCC |
| | qPCR_ets-4 F span | AAAGACAACGACGTGTTGCTATCTG |
| | qPCR_ets-4 R span | GACACAATAGGAATATGTTCTACAACG |
| | tbb-2 qPCR R | TGGTGAGGGATACAAGATGG |
| | tbb-2 qPCR F | GCTCATTCTCGGTTGTACCA |
| | GFP F | GTTGTCCCAATTCTTGTTGAATTAGATGG |
| | GFP R | TCGAGAAGCATTGAACACCA |
| *unc-54* amplification for restriction enzyme cloning | unc-54 F XhoI | CTAGCTCGAGGTCCAATTACTCTTCAACATCC |
| | unc-54 R NotI | CTAGGCGGCCGCCAAAAAAATTTATCAGAAGTAAAAAAC |
| *Il6* amplification for restriction enzyme cloning | Il6 403 F XhoI | CTAGCTCGAGTGCGTTATGCCTAAGCATATCAG |
| | Il6_NotI 1-403 R | CTAGGCGGCCGCTTTGTTTGAAGACAGTCTAAAC |
| *ets-4* amplification for restriction enzyme cloning | ets-4 F XhoI | CTAGCTCGAGTCATCTGGCAGAAAGACAACGACGT |
| | ets-4 R NotI | CTAGGCGGCCGCAGATTATGAGACCTTTGGACTTG |
| F1S amplification for Gibson cloning | F1S F gibson | TTCTCTTAATTTCTTTGTGGTCAATACTCTGTTTACATTTTTC |
| | F1S R gibson | AAAGAAGCTAAAAAGGCGCGAGGAATATGTTCTACAACG |
| *Ox40* amplification for restriction enzyme cloning | Ox40 F | CTAGCTCGAGGCATTACTAC |
| | Ox40 R | CTAGGCGGCCGCGCCAGTC |
| REGE-1 cDNA amplification for restriction enzyme cloning | REGE-1 F NotI | CTAGGCGGCCGCATGGATTCAACGGCTCGTGG |
| | REGE-1 R KpnI | GGTACCCTAG |
| Overlapping primers to create F1SΔRCE fragment | F1SdRCE F | TTCTCTTAATTTCTTTGTGGTCAATACTCTGTTTACATTTTTCAACTTGTCCTGTCGTTCAAGATATAC |
| | F1SdRCE R | AAAGAAGCTAAAAAGGCGCGAGGAATATGTTCTACAACGAACAGTACATGGAAGTATATCTTGAACGACAGGAC |

| | | |
|---|---|---|
| Amplification of RLE-1 cDNA with tags | RLE-1 amp F pCS2 | ATGGACTACAAAGACGATGACGACAAGATGGCGCCAAC GGGTCAAGGTGGGC |
| | RLE-1 amp R pCS2 | TCAcagatcctcttcagagatgagtttctgttcTCCCTCGACAGTCGGA TTG AGATGAAGC |
| Amplification of REGE-1 cDNA with tags | REGE-1 amp F pCS2 | ATGGACTACAAAGACGATGACGACAAGATGGATTCAAC GGCTCGTGGCCAC |
| | REGE-1 amp R pCS2 | TCAcagatcctcttcagagatgagtttctgttcTTTTCGGTACTCTTTTT GAGCTCGGATAATG |
| Amplification of N-FLAG, C-MYC sequence for Gibson cloning | FLAG gibs pCS2 F | CTACTTGTTCTTTTTGCAGGATCCCATATGGACTACAAA GACGATGACGACAAG |
| | MYC gibs pCS2 | CTACGTAATACGACTCACTATAGTTTCAcagatcctcttcagag atgagtttctgttc |
| Amplification of templates for *in vitro* transcription | F1S F T7 | TAATACGACTCACTATAGGGTCAATACTCTGTTTACATT T |
| | F1S R | AGGAATATGTTCTACAACGAACAG |
| | F1SdADE R | AGGAATATGTTCTACAAGGAATAG |
| | F1SdREGE R | AGGAATATGTTCTACAACGAACAG |
| SHAPE | ets-4 SHAPE | AGGAATATGTTCTACAACGAACAGT |

All oligonucleotides used in this study were ordered from Merck, Germany.

**SUPLEMMENTARY REFERENCES**


1. Hermann, G.J., Schroeder, L.K., Hieb, C.A., Kershner, A.M., Rabbitts, B.M., Fonarev, P., Grant, B.D. and Priess, J.R. (2005) Genetic analysis of lysosomal trafficking in Caenorhabditis elegans. *Mol Biol Cell*, **16**, 3273-3288.
